# Molecular dynamics simulation of local structural models of PrP^Sc^ reveals how codon 129 polymorphism affects propagation of PrP^Sc^

**DOI:** 10.1101/2019.12.25.888289

**Authors:** Hiroki Otaki, Yuzuru Taguchi, Noriyuki Nishida

**Author notes:** The two authors equally contributed to the work (Hiroki Otaki, Yuzuru Taguchi).

## Abstract

Prions are unconventional pathogen without nucleotide genome and their pathogenic properties are defined by the primary structure and the conformation of the constituent abnormal isoform (PrP^Sc^) of prion protein (PrP). A polymorphic codon 129 of human PrP that is valine (V129) or methionine (M129) is particularly influential on properties of PrP^Sc^, affecting transmission efficiencies and clinicopathological features. However, how the single residue is so influential has not been elucidated because the detailed structures of PrP^Sc^ have not been determined yet due to its incompatibility with high-resolution structural analysis. Previously we created an in-register parallel β-sheet local structural model of human PrP^Sc^ encompassing residues 107 to 143 that seemed more compatible with V129 than M129, based on knowledge from α-synuclein amyloids and an NMR-based model of the amyloid of Y145Stop mutant of PrP in the literature. Here, we created an M129-compatible local structural model of PrP^Sc^. Severe destabilization of the model by G127V mutation was consistent with the protective effects of V127 polymorphism of human PrP against prions. It was highly sensitive to the length of the hydrophobic side chain of codon 129 and replacement of M129 with leucine or valine destabilized the structures. Interestingly, the U-shaped β-arch which comprises M129 flexibly changed hydrophobic interaction networks inside the β-arch depending on the interactions with the surrounding structures, whereas the previous model with V129 maintained the similar network patterns irrespective of the surroundings. The differences between the two models may explain influences of the codon 129 polymorphism on transmissions and properties of human prions.

## Introduction

Prions are unconventional pathogens composed solely of prion protein (PrP) without any nucleotide genome. Prions cause diseases characterized by accumulation of the abnormal isoform of PrP (PrP^Sc^) in the central nervous system [1], and they can occur in various mammalian species because many species constitutively express PrP, e.g., Creutzfeldt-Jakob disease (CJD) in humans, bovine spongiform encephalopathy (BSE) in cattle [2], and chronic wasting disease (CWD) in cervids [3]. Despite the lack of nucleotide, prions show virus-like properties, including high infectivity, existence of many prion strains, and species/strain barriers. As the virus-like pathogenic properties are encoded in the conformations of PrP^Sc^ [1][4], the pathogenic phenotypes greatly depend on the primary structure of the constituent PrP^Sc^. Most demonstrative examples of such phenomena are the influences of polymorphic codon 129 (in human numbering unless otherwise noted) of human PrP, which is either methionine (M129) or valine (V129). Clinicopathological features of sporadic CJD vary depending on the polymorphism, in terms of PrP deposition patterns and lesion profiles in the brain, and apparent molecular sizes of protease-resistant cores of PrP^Sc^ [5]. The PrP^Sc^ from CJD patients homozygous for M129 tend to form diffuse tiny deposits in the brain with 21-kDa protease-resistant core fragment. On the other hand, PrP^Sc^ from CJD heterozygous or homozygous for V129 often forms plaque-type and perineuronal deposits with 19-kDa protease-resistant core [5]. The codon 129 polymorphism also affects transmission efficiencies of prions. For example, a vast majority of patients with new-variant CJD, manifestation of trans-species transmission of BSE to humans, are homozygous for PrP with M129 (M129-PrP) [6][7]. In transmission experiments of CJD prions to transgenic (Tg) mice expressing human-mouse chimeric PrP that has mouse sequences only in the octapeptide-repeat region and the distal part of the third α-helix (H3) [8], incubation periods of Tg mice expressing chimeric M129-PrP were significantly shortened, whereas Tg mice expressing chimeric V129-PrP were resistant to the transmission. The methionine/leucine polymorphic codon 132 (in elk numbering) of elk PrP equivalent to the codon 129 of human PrP also affects susceptibility to CWD [9][10]. The molecular mechanism of how differences between the hydrophobic amino acids greatly affects properties of PrP^Sc^ is intriguing but extremely challenging. Although detailed structures of PrP^Sc^ are essential for the investigation, they are currently unavailable due to the difficulty in high-resolution structural analyses of PrP^Sc^. Even whether PrP^Sc^ is an in-register parallel β-sheet amyloid or a β-solenoid is still controversial [11][12][13][14][15][16]. Regardless, we previously created a local structural model of PrP^Sc^ encompassing residues 107 to 143 [17] by assuming that PrP^Sc^ is an in-register parallel β-sheet amyloid and designing based on a structural model of Y145Stop-mutant PrP amyloid reported by Theint et al. [16][18]. The model seemed structurally more stable, i.e., compatible, with V129 than M129 [17].

Then, we attempted to create models that are more compatible with M129 than V129 or L129, because PrP of the majority of humans and most species have M129-PrP. Barriers in transmissions of CJD or CWD from M129-PrP-expressing individuals to those with V129- or L129-PrP, respectively, suggest that codon 129 defines the preference of the substrate PrP^C^ for the conformation of PrP^Sc^. Therefore, conformations of M129-compatible PrP^Sc^ and V129-compatible PrP^Sc^ should be sufficiently different accordingly. Interestingly, the traits of V129-compatible PrP^Sc^ are so stable that they are preserved through passages to M129-PrP-expressing individuals, without abbreviation of incubation periods on the secondary transmissions, i.e., no adaptation to M129-PrP [19][20][21], reminiscent of non-adaptive prion amplification, NAPA [22]. This was in contrast with ordinary species barriers accompanied by adaptation to the new host, e.g., transmission of Syrian hamster scrapie to Chinese hamsters [23]. Possibly, the conformations of M129- and V129-compatible PrP^Sc^ are not readily interchangeable. On the other hand, they may share certain similarities: protease-resistant core of mouse-adapted scrapie 22L prion, whose protease-cleavage sites should be N-terminal to the residue 89 like 21-kDa core [24], was tolerant to insertion of a tetracysteine tag up to the residue 96 [25], which corresponds to the N-terminus of the 19-kDa core [26]. Another mouse-adapted scrapie also tolerated insertion of a linear peptide up to 94 [27]. Those findings suggested that the compactly folded moiety of the 21-kDa core that is preferred by M129 may be similar to that of the 19-kDa core preferred by V129, and the more N-terminal part of the former is just loosely folded. What conformations could fulfill those requirements? If M129- and V129-compatible PrP^Sc^ are remotely different as tau amyloids from Alzheimer’s and Pick’s diseases [28], they would not be interchangeable but have the less similarity.

Here, we demonstrate that even a relatively small structural differences may suffice. Our new model was still based on the Theint’s model, but a β-sheet encompassing residues 120-122, β-sheet(120-122), was flipped and only valine at 121 (V121) faced the side chain of M129 (**Fig 1A**). In molecular dynamics (MD) simulations, mutations equivalent to polymorphisms that affect transmission efficiencies of prions with M129, e.g., G127V, M129V, and M129L, substantially affected stability of the amyloids. The present model provides an example of local structures of PrP^Sc^ whose stability is highly sensitive to the length of hydrophobic side chain at codon 129. Moreover, truncated mutants of the model provided insights into the mechanism of conformation determination of PrP^Sc^ depending on M/V129 polymorphism [29]. The explanation of effects of the polymorphisms by our models further supports the view that PrP^Sc^ is an in-register parallel β-sheet amyloid. Those knowledge from the present studies would advance our understanding about strain diversity of amyloids, and also aid in designing more plausible PrP^Sc^ models and other amyloids in the future.

**Fig 1.**
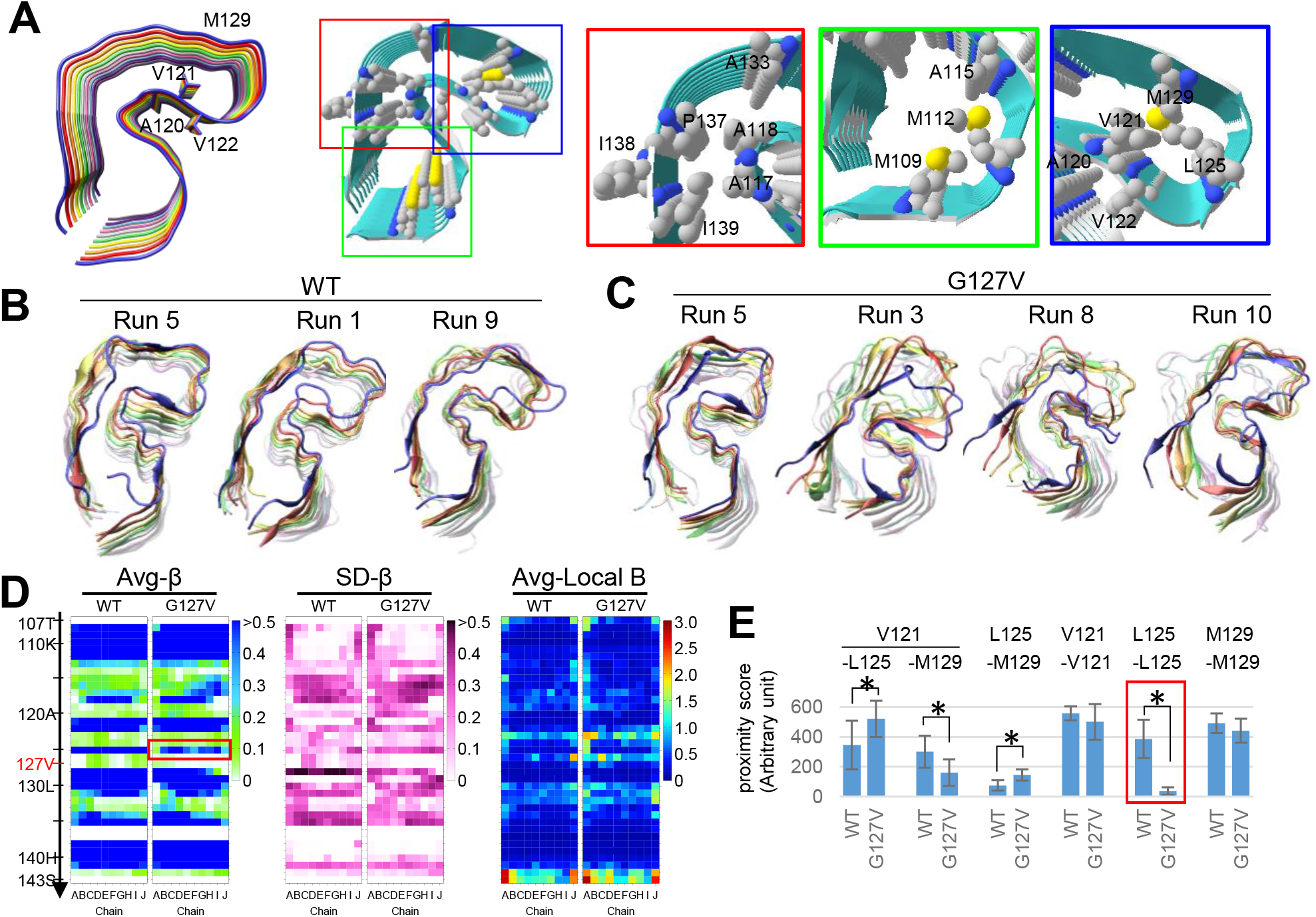
Designing a local structural model of human PrP^Sc^ encompassing residues 107-143 and assessment of influences of G127V mutation on the structural stability. **A**. (Left) A PrP(107-143) amyloid as a tentative local-structural model of PrP^Sc^, which consists of 10 layers (chains A–J) like the previous model **[17**]. Note that V121 is facing M129 in this model. (Right) Side chains of the hydrophobic residues whose hydrophobic contacts are important for structures are explicitly exhibited (red~blue insets). **B-C**. Final snapshots of (**B**) the wild-type (WT) and (**C**) the G127V-mutant after a 10-ns MD simulation. A relatively stable one (left) and unstable ones (middle and right) of ten independent runs of MD simulation are selected: final shots of all the ten runs are presented in **Supplementary Figure S1**. **D**. (Left and middle) Heatmaps showing the average values and the standard deviation values of β-sheet propensity of each residue (Avg-β and SD-β, respectively) of the WT and the G127V-mutant PrP amyloids (WT and G127V). The values were calculated based on ten independent runs of 10-ns MD simulation. Note that the Avg-β values are lower in the G127V in the row of L125 (red box). (Right) Heat maps of local backbone B-factors of each residue, which is a metric for local backbone flexibility, of WT and G127V amyloids. **E**. A Graph of the proximity scores for representative hydrophobic interactions that may contribute to stability of the U-shaped β-arch. The bars and error bars represent mean ± standard deviation (SD) of values from ten independent runs. The asterisks indicate p < 0.05. Note that many interactions are significantly affected by the G127V mutation and particularly L125-L125 interactions are drastically impaired (red box).

## Results and Discussion

### Configuration of a local structural model of M129-compatible PrP^Sc^

#### MD simulation of Wild-type (WT) model with M129

The protective effects of L132 of elk PrP against CWD (in elk numbering) [9][10] and transmission barriers between M129-PrP- and V129-PrP-expressing individuals [19][30] implied that length of the hydrophobic side chain of codon 129 is critically important for propagation of M129-compatible PrP^Sc^. We already had an idea about what structure could exhibit such properties through MD simulation of G84M- and G84I-mutants of α-synuclein amyloids [17]. Therefore, we created a new model by flipping β-sheet(120-122) of the previous model (**Fig 1A, left**) so that the side chain of M129 barely maintain contact with that of V121 in the hydrophobic core of the U-shaped β-arch encompassing 120 to 133. We also modified the surrounding structures by arbitrarily positioning hydrophobic residues to let them efficiently interact and stabilize the local structures (**Fig 1A, right panels**). In the MD simulations, the model was sufficiently stable for analyses (**Fig 1B, left**). Even in relatively disordered runs, chain A or J only partly dislocated from the stack at the U-shaped β-arch (**Fig 1B, middle and right; Suppl Fig S1, WT**). In the cases where we compare the present model with the previous model [17], to avoid confusion, we refer to the present and the previous models as ‘Type-1’ and ‘Type-2’ models/β-arches, respectively, based on the numbers of hydrophobic residues mainly interacting with residue 129 (i.e., only V121 in Type-1, while A120 and V122 in Type-2) (**Suppl Fig S2A**).

#### MD simulation of the G127V mutant

First, we observed influences of G127V substitution that corresponds to a well-known protective polymorphism of human PrP against CJD [31] on structural stability of the model. The G127V amyloid obviously appeared unstable and the destabilization was so extensive that the chains were unraveled in the U-shaped β-arch (**Fig 1C**) as in the previous/Type-2 model [17]. Heat maps of β-sheet propensity values confirmed the structural instability (**Fig 1D, G127V**). The average β-sheet propensity values were particularly lower at L125 compared with WT, confirming the unraveling (**Fig 1D, red box**). The lower β-sheet propensities at L125 was consistent with the significantly lower proximity scores for L125-L125 interactions than that of WT (**Fig 1E, red box**), which indicates persistence of hydrophobic contacts between the two residues [17]. As in-register parallel β-sheet amyloids have ~4.8 Å intervals between the chains, the proximity score with cut-off value of 5 Å well reflect the stability of the β-sheets. Besides L125-L125 interactions, other hydrophobic interactions in the hydrophobic core of the β-arch were also significantly affected, e.g., V121-M129 (**Fig 1E**). The heat map of local B-factors of the G127V amyloid (**Fig 1D, local B**), which is a metric for local flexibility of backbone [32], showed more bright spots in the region of the U-shaped β-arch(120-133) than that of WT, further confirming the structural disorder. Interestingly, the hydrophobic interaction network outside the U-shaped β-arch was similar to that of WT (**Suppl Fig S2B, G127V**). The destabilization by G127V substitution was consistent with its protective effects against CJD prions.

#### MD simulation of the M129V mutant

Then, we tested our initial assumption that M129 is more advantageous to maintain the hydrophobic core than V129 or L129 by observing influences of M129V or M129L substitutions. The M129V amyloid seemed less stable than WT (**Fig 2A**) with more frequent loss of β-sheet structures and dislocation of chain A or J from the stack (**Suppl Fig S1, M129V**). The heat maps of average β-sheet propensities confirmed the unstable β-sheets of the M129V mutant, particularly at L125 and in β-sheet(128-130) (**Fig 2B, inset in red)**. The heat map of local B-factors of the M129V amyloid was apparently more spangled with bright spots than that of WT, particularly at G126 **(Fig 2B, green inset)**. Proximity scores of the hydrophobic interactions in the hydrophobic core of the U-shaped β-arch provided more quantitative differences between WT and M129V. The most notable finding was the significantly lower scores for L125-L125 interactions of M129V than those of WT (**Fig 2C, blue box**). Although not as much as G127V, the impaired interactions confirmed the structural instability of the M129V amyloid. Likewise, proximity scores for V129-V129 interactions of M129V were also lower than M129-M129 of WT, though not statistically significant (490.66 ± 65.50 vs 424.98 ± 80.78, p = 0.075). This is in contrast with the results of the previous/Type-2 model, where the scores for the V129-V129 interactions of M129V mutant tended to be higher than those of the WT (449.15 ± 111.65 vs 513.31 ± 62.17; not statistically significant) [17]. Unexpectedly, the proximity scores for V121-V129 of M129V and WT were comparable despite the short length of the side chain of V129 (**Fig 2C**). Accordingly, the distances between Cα of V121 and V129 tended to be shorter than V121-M129 of WT (**Fig 2E, M129V**). The high intrinsic β-sheet tendency of V129 may somehow bring β-sheet(120-122) and β-sheet(128-133) closer, as most prominently demonstrated in Run 10, where the two β-sheets approached so close as to form a triangular conformation reminiscent of the M129V mutant of the previous/Type-2 model [17] (**Suppl Fig S1, M129V**). Unlike the previous/Type-2 model, the closer positioning of the β-sheets seemed to cost the structural integrities in the present/Type-1 model. The relatively higher proximity scores for V121-L125 interactions of M129V (**Fig 2C**) may reflect altered positional relations of V121 and L125 due to the distortion of the β-arch.

**Fig 2.**
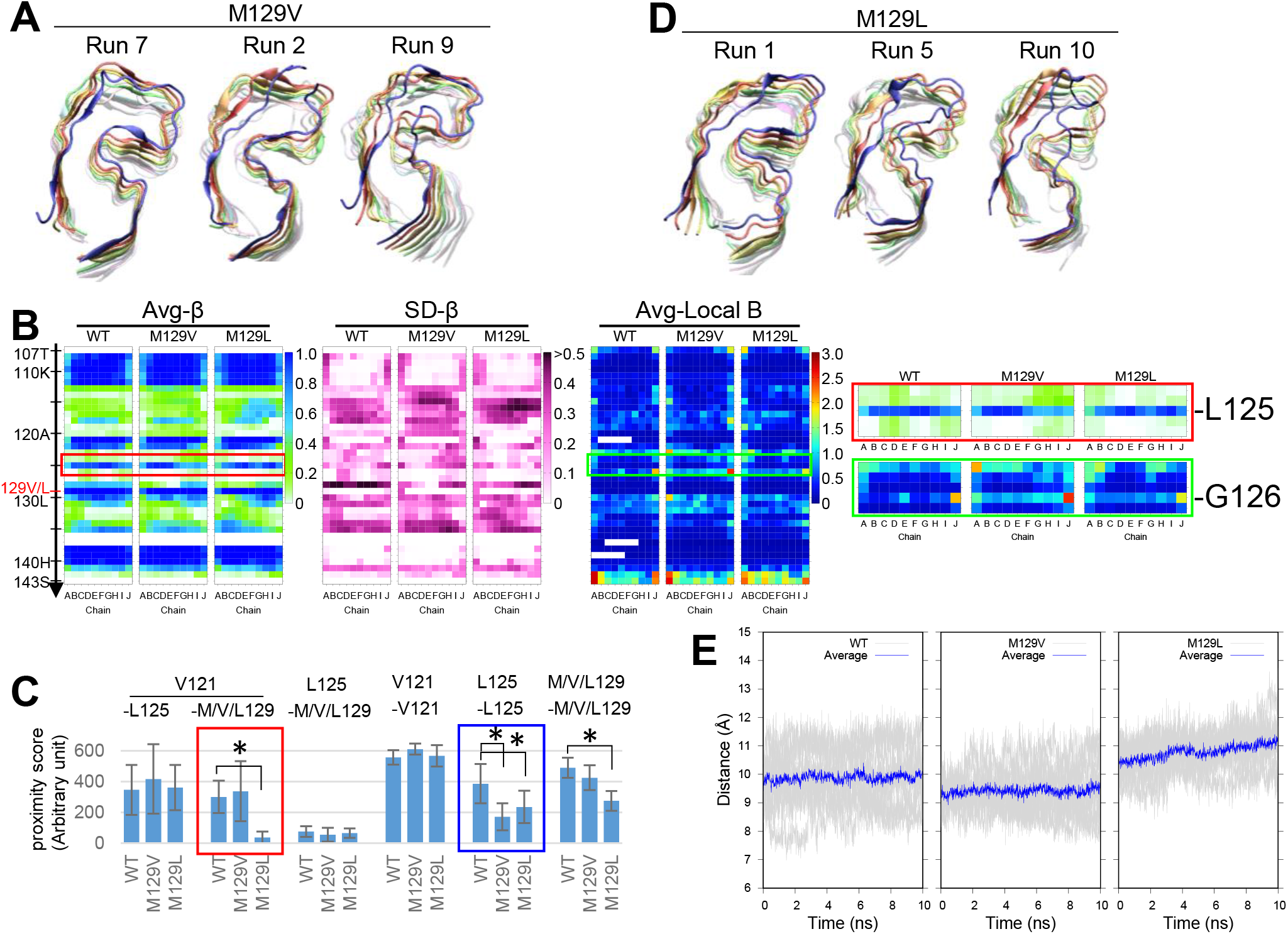
M129V or M129L substitutions destabilize the amyloid structures. **A**. Final snapshots of M129V-mutant amyloid after a 10-ns MD simulation. A relatively stable one (left) and unstable ones (middle and right) of ten independent runs of MD simulation are selected: final shots of all the ten runs are presented in **Supplementary Figure S1**. **B**. showing Avg-β (left) and SD-β (middle) values of each residue of the WT and the mutant amyloids (WT, M129V and M129L). (Right) Heat maps of average local B-factors of each residue. (Red and green insets) Magnified views of the Avg-β and local backbone B-factors around L125 residue, respectively. The values were calculated based on ten independent runs of 10-ns MD simulation. Note that the Avg-β values of the L125 row are lower (paler) in M129V and M129L compared with that of WT, suggesting instability in β-sheet structures in the mutant. **C**. A Graph of proximity scores for hydrophobic interactions that may contribute to stability of the U-shaped β-arch, comparing WT, M129V and M129L amyloids. The bars and error bars represent mean ± SD of values from ten independent runs. The asterisks indicate p < 0.05. Note that V121-L129 and L125-L125 interactions are significantly lower in the M129L amyloid (red and blue boxes, respectively). **D**. Final snapshots of PrP(107–143;M129L)-mutant amyloid after a 10-ns MD simulation. A relatively stable one (left) and unstable ones (middle and right) of ten independent runs of MD simulation are selected. Final shots of all the ten runs are presented in **Supplementary Figure S1**. **E**. Graphs showing distances between Cα atoms of V121 and M/V/L129 of chain E (a chain at the middle of the stack) over 10 ns of MD simulations. The gray lines represent the values of each run of ten independent simulations, and the blue line represents the average values.

#### MD simulation of the M129L mutant

The M129L amyloid also appeared more unstable than the WT, with chain A more frequently dislocating from the stack (**Fig 2D**). The heat maps of β-sheet propensities and local B-factors exhibited similar changes as those of M129V but of milder degrees (**Fig 2B, M129L**). In proximity score analysis, significantly lower L125-L125 interactions than that of WT also corroborated the instability (**Fig 2C, blue box**). The most remarkable finding was the drastically low scores for V121-L129 interactions (**Fig 2C, red box**). The wider distances between Cα of V121 and L129 was consistent with loss of the bond between them (**Fig 2E, M129L**), which could be attributed to the shorter side chain of L129 than M129. In addition, the significantly lower scores for L129-L129 interactions suggested attenuated inter-layer interactions of β-sheet(128-133) (**Fig 2C**). The lack of Cβ-branching of L129 may be responsible for the different effects from those of V129, because it allows the back bone more freedom of conformation.

Those findings confirm the instability of M129V and M129L amyloids relative to WT, supporting our view that length of the hydrophobic side chain of codon 129 is critical for the stability of the present/Type-1 model.

#### MD simulation of the I138M mutant

In the previous model of the local structure of PrP^Sc^ [17], methionine substitution at the residue 138 that is different between human and mouse (Ile and Met, respectively) caused substantial alterations in the conformation of the amyloid. We therefore assessed influences of I138M mutation on the present model. The final shots of I138M amyloid seemed more stable than WT, without any dislocation of chain A from the stack in ten independent runs of simulation (**Fig 3A**). β-sheet propensities and local B-factors were also comparable to those of WT (**Fig 3B**). Indeed, the average proximity score for L125-L125 interactions was higher than that of WT (**Fig 3C**), whereas the scores for V121-M129 interactions were significantly lower (**Fig 3C, red box**). This could be due to the wider distances between the Cα of V121 and M129 (**Fig 3D**). The widening may be secondary to the structural alteration in the C-terminal region by the I138M substitution that makes the local back-bone more flexible, although specifically what alteration was not discernible [17]. This mutant suggests that V121-M129 interactions does not have to be very high for stability of the U-shaped β-arch.

**Fig 3.**
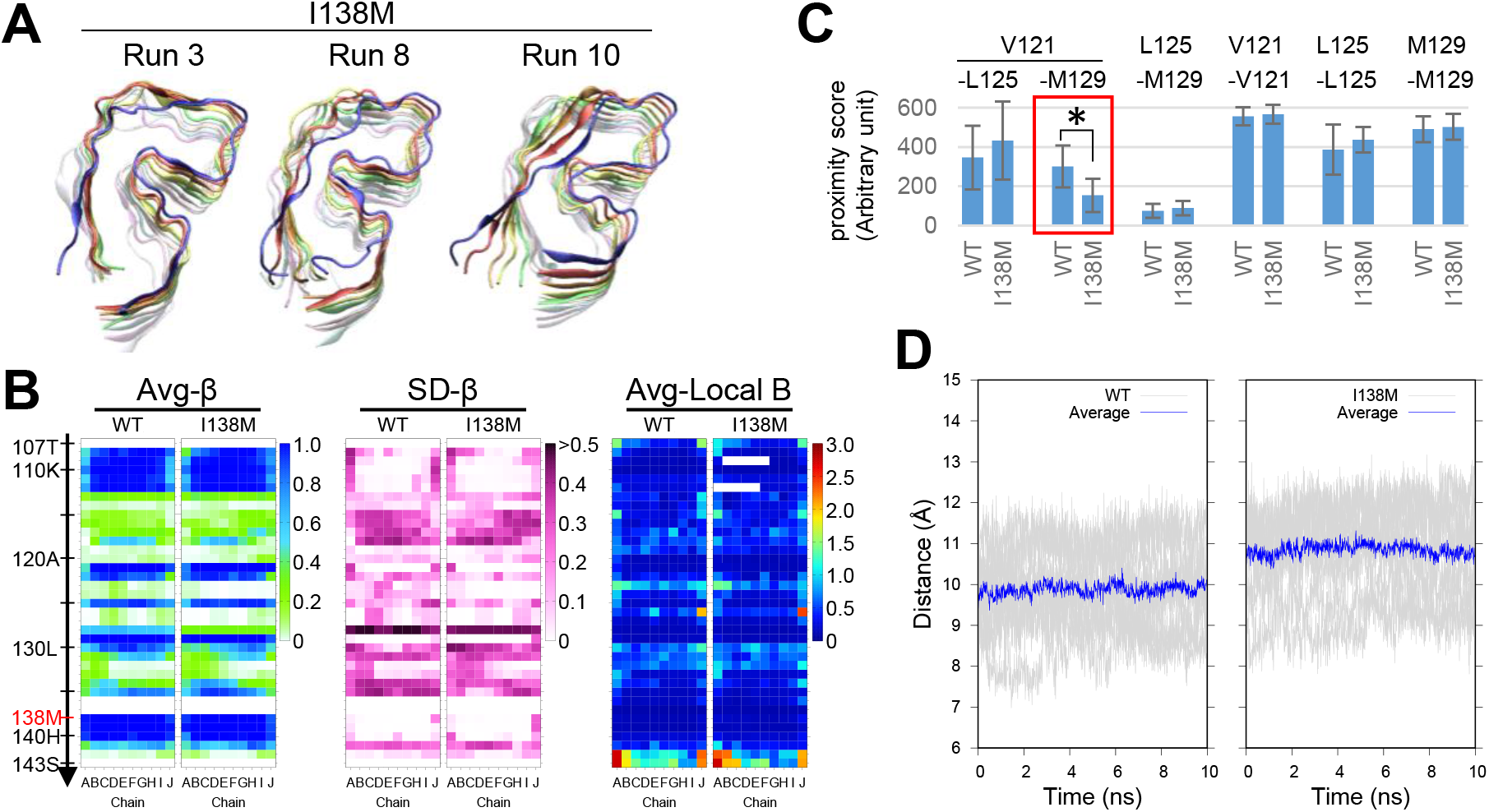
I138M mildly diminished V121-M129 interactions by widening the loop but did not affect the structural stability of the amyloid. **A**. Final snapshots of the I138M mutant amyloid after a 10-ns MD simulation. A relatively stable one (left) and unstable ones (middle and right) of ten independent runs of MD simulation are selected: final shots of all the ten runs are presented in **Supplementary Figure S1**. **B**. (Left and middle) Heat maps showing Avg-β and SD-β values of each residue of WT and I138M-mutant amyloids (WT and I138M). (Right) Heat maps of local backbone B-factors. The values were calculated based on ten independent runs of 10-ns MD simulation. **C**. A graph of proximity scores for hydrophobic interactions that may contribute to stability of the U-shaped β-arch, comparing WT and I138M amyloids. The bars and error bars represent mean ± SD of values from 10 independent simulation runs. The asterisks indicate p < 0.05. Note that the V121-M129 interactions are significantly lower in the I138M mutant amyloid (red box). **D**. Graphs showing distances between Cα atoms of V121 and M129 of chain E over 10 ns of MD simulations. The gray lines represent the values of each run of ten independent simulations, and the blue line represents the average values.

#### Stability of the amyloids of truncated peptides in the present/Type-1 conformation

The interaction network diagram of the Type-1 model showed multiple intense interactions between the U-shaped β-arch and the surrounding structures, e.g., A118-P137 and A117-I139 (**Suppl Fig S2B**). To evaluate their contribution to the stability of the U-shaped β-arch, we created amyloids of the C-terminally truncated molecules, PrP(107-137) of the Type-1 model (C-truncated WT) and its M129V mutant, and observed on MD simulation. Unexpectedly, the C-terminal truncation of WT further stabilized the U-shaped β-arch (**Fig 4A**), while accentuating the instability of M129V mutant (**Fig 4B**). Heat maps of β-sheet propensity and local B-factors confirmed less β-sheet structures and the instability in the U-shaped β-arch (**Fig 4C, red boxes**). Interaction network diagrams confirmed the loss of the intense A118-P137 and A117-I139 interactions in both the amyloids but, instead C-truncated WT reinforced the U-shaped β-arch by newly forming intense A118-A133 and A113-V122 interactions that were not Type-1 before truncation (**Fig 4D, WT**). In contrast, C-truncated M129V also formed A118-A133 but the proximity score was ~5-fold lower (**Fig 4E, blue box**), and the network in the N-terminal region was quite similar with that of the full-length WT (**Fig 4D, M129V**). Interactions in the hydrophobic cores of the U-shaped β-arches were also rather different (**Fig 4E, bold letters**); WT showed nearly even scores for the three interactions albeit slightly higher V121-L125, whereas M129V had outstanding V121-V129 interactions ~5-fold higher than others. As anticipated, the differences in the scores of L125-L125 interactions between the truncated molecules were also accentuated by the truncation (**Fig 4E, red box**).

**Fig 4.**
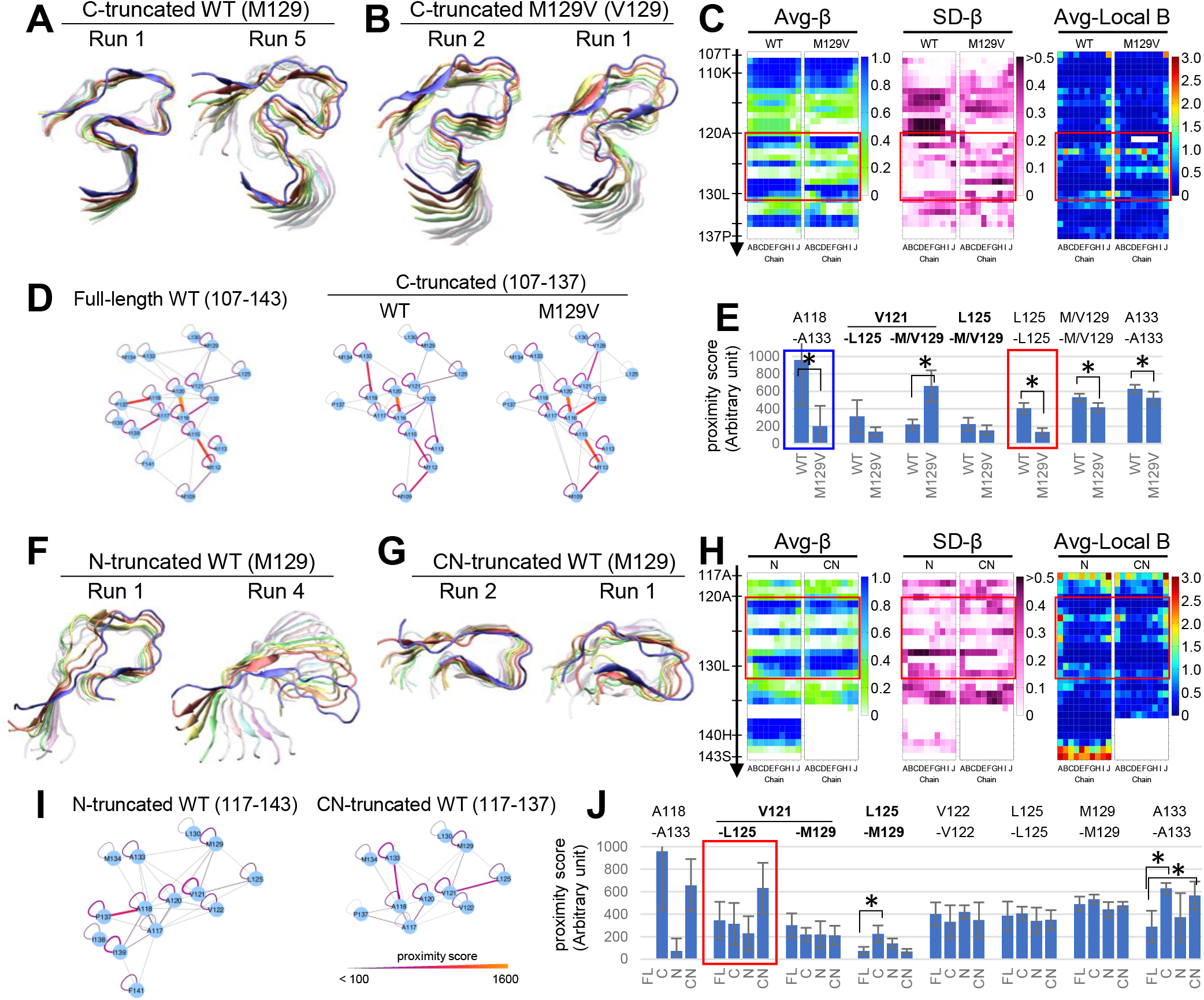
MD simulation of truncated mutants of the present local structural model of PrP^Sc^ revealed its flexibility. **A-B**. Final shots after 10-ns MD simulations of (**A**) the C-terminally truncated models (encompassing 107-137) of the wild-type (WT) and (**B**) the C-terminally truncated M129V mutant (M129V). A relatively stable (left) and a relatively unstable ones (right) are selected from five independent runs. Final shots of all the five runs are presented in the **Supplementary Figure S3**. **C**. Heat maps of Avg-β (left), SD-β (middle) and average local backbone B-factors (right) of the C-truncated WT and M129V. The values are based on five independent runs of 10-ns simulations. Note that Avg-β values in the U-shaped β-arch region, particularly in the row of L125, are substantially decreased, SD-β values are increased and Avg local backbone B-factors are increased in M129V (red boxes). **D**. Hydrophobic interaction diagrams of the full-length WT (left), the C-truncated WT (middle) and M129V (right). The color and thickness of the lines represent the proximity scores of the interactions and the legend is presented in Fig I. **E**. A graph of proximity scores for hydrophobic interactions comparing the C-truncated WT and C-truncated M129V. The interactions in bold letters indicate those in the hydrophobic core of the U-shaped β-arch. The bars and error bars represent mean ± SD of scores from five independent runs of 10-ns simulation. Note that L125-L125 interactions are significantly lower in the mutant than that of WT (red box). Asterisks indicate p < 0.05. **F-G**. Final shots after 10-ns MD simulations of (**F**) the N-terminally truncated models (encompassing 117-143) and (**G**) the C-terminally truncated model (117-137) of the WT amyloid. A relatively stable (left) and a relatively unstable ones (right) were selected. Final shots of all the five runs are presented in the **Supplementary Figure S3**. **H**. Heat maps of Avg-β (left), SD-β (middle) and average local backbone B-factors (right) of the N-truncated (N) and the double-truncated (CN) amyloids. The values are based on five independent runs of 10-ns simulations. Note that Avg-β, SD-β and Avg local backbone B-factors are rather similar in the U-shaped β-arch region between the amyloids (red boxes). **I**. Hydrophobic interaction diagrams of the N-truncated (left) and the CN-truncated WT (right). **J**. A graph of proximity scores for hydrophobic interactions comparing the full-length WT (FL), C-truncated WT (C), N-truncated WT (N) and double-truncated WT (CN). The bars and error bars represent mean ± SD of scores from five (ten for FL) independent runs of 10-ns simulation.

We created amyloids of further-truncated molecules of the WT molecule, N-terminally (N) truncated WT PrP(117-143) and C- and N-terminally (CN) truncated WT PrP(117-137), assuming that the PrP^C^-PrP^Sc^ conversion process initiates by incorporating the native β-sheets of the substrate PrP^C^, β1 (encompassing 128-131), into the template PrP^Sc^ and the truncated amyloids provide insights into the initial phase of the conversion. The N-truncated and the CN-truncated amyloids appeared comparably unstable (**Fig 4F and G**), suggesting the C-terminal truncation of the N-truncated amyloid did not further destabilize. Heat maps confirmed their similar stabilities, despite the differences in their sizes (**Fig 4H**). Interaction network diagrams of the N-truncated amyloid was reminiscent of that of the full-length amyloid due to intense A118-P137 and very weak A118-A133 interactions and the similar pattern in the hydrophobic core (**Fig 4I**). On the other hand, CN-truncated amyloid resumed intense A118-A133 like the C-truncated, but interactions in the hydrophobic core were predominated by outstandingly intense V121-L125 interactions (**Fig 4J, red box**). Thus, each truncated model showed its unique pattern of interaction network to maintain stability, as indicated by proximity scores of L125-L125 interactions (**Fig 4J**).

#### Stability of the amyloids of truncated peptides in the previous/Type-2 conformation

The results of the truncated models prompted us to MD simulation of the C- or/and N-terminal truncated amyloids of the previous model [17]. As stated earlier, we refer to the previous model as ‘Type-2’ model/β-arch.

We used Type-2 model with V129 for the truncation and MD simulation, because the M129V mutant was more stable than WT. In contrast to the Type-1 model, C-terminal truncation substantially destabilized the amyloid structures (**Fig 5A**), and N- and CN-truncations additively further destabilized (**Fig 5B and C**). Heat maps confirmed the gradual decrements in β-sheet propensities in the β-arch region (**Fig 5D, red box**). A remarkable finding was in their interaction diagrams: they showed very similar patterns of interaction networks irrespective of truncation types (**Fig 5E**), albeit modest alterations in intensities of some bonds (**Fig 5F**). That was reminiscent of the C-truncated V129 Type-1 model (**Fig 4D, M129V**). The significant decrements in the L125-L125 interactions confirmed the aggravation of instability by the truncations (**Fig 5F, red box**).

**Fig 5.**
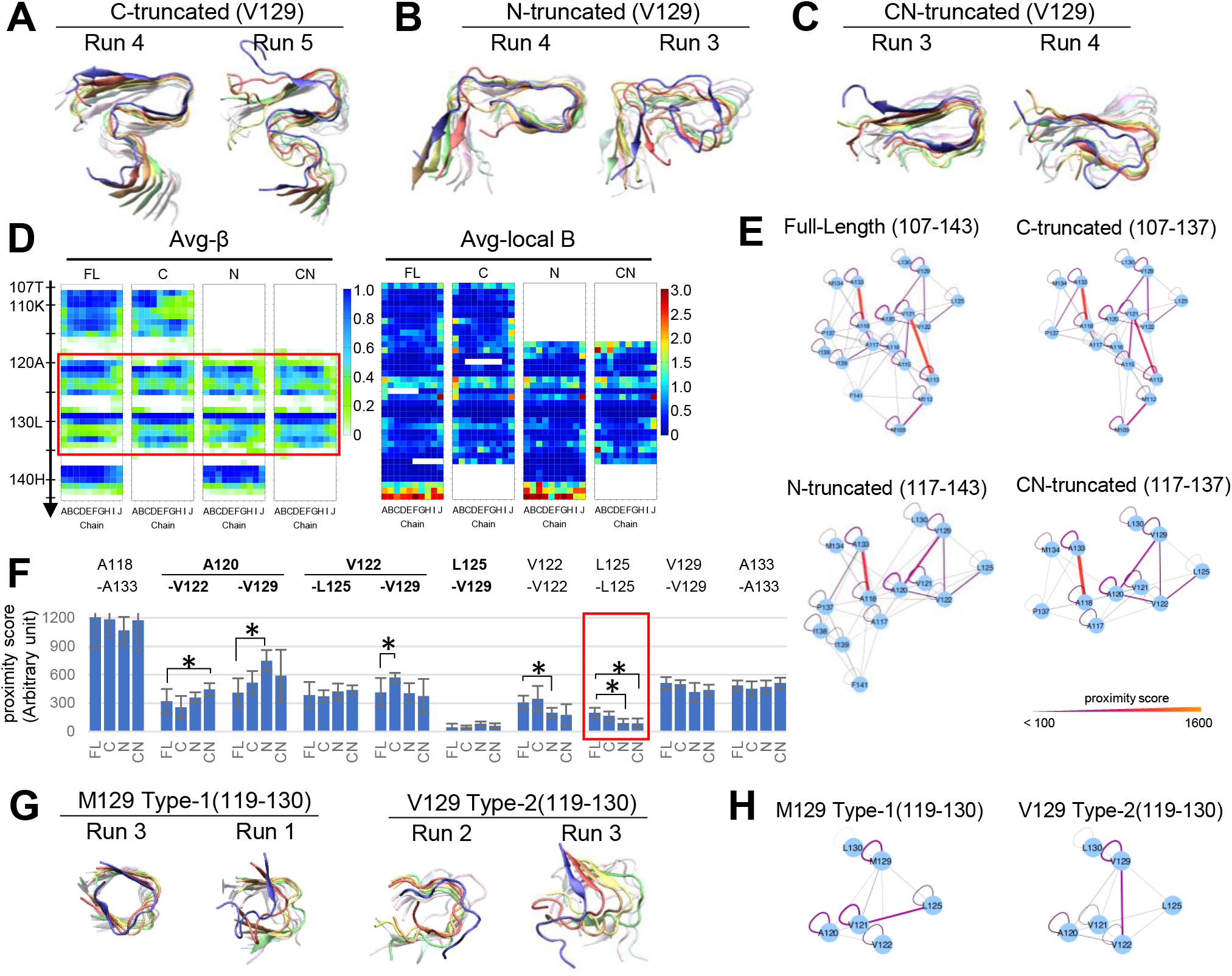
MD simulation of truncated mutants of the previous local structural model of PrP^Sc^ (V129 Type-2 model) revealed its rigidity. **A-C**. Final shots after 10-ns MD simulations of (**A**) C-terminally truncated V129 Type-2 model encompassing 107-137, (**B**) N-terminally truncated model encompassing 117-143, and (**C**) C- and N-terminally truncated model encompassing 117-137. A relatively stable (left) and a relatively unstable ones (right) are presented. Final shots of all the five runs are presented in the **Supplementary Figure S3**. **D**. Heat maps of Avg-β (left) and average local backbone B-factors (right) of the full-length (FL), C-truncated (C), N-truncated (N) and CN-truncated (CN). The values are based on five independent runs of 10-ns simulations. Note that Avg-β values in the U-shaped β-arch region (red box) are substantially decreased in the truncated amyloids. **E**. Hydrophobic interaction diagrams of the full-length and the truncated V129 Type-2 models. The color and thickness of the lines represent the proximity scores for the interactions. Note that the interaction networks in the U-shaped β-arches are rather similar irrespective of the truncation types. **F**. A graph of proximity scores for representative hydrophobic interactions. The interactions in bold letters indicate those in the hydrophobic core of the U-shaped β-arch. The bars and error bars represent mean ± SD of scores from five (ten for FL) independent runs of 10-ns simulation. Note that L125-L125 interactions are significantly lower in the N- or CN-truncated amyloid than that of the full-length (red box). Asterisks indicate p < 0.05. **G**. Final shots after 10-ns MD simulations of peptides corresponding to the residues 119-130 of M129 Type-1 model and V129 Type-2 models. A relatively stable (left) and a relatively unstable ones (right) are presented. Final shots of all the five runs are presented in the **Supplementary Figure S4**. H. Hydrophobic interaction diagrams of the full-length and the peptides 119-130. The color and thickness of the lines represent the proximity scores for the interactions.

#### Stability of the further-truncated Type-1 and Type-2 amyloids

We further truncated the amyloids, i.e., encompassing G119 to G131, so that the A117-A133 interactions are eliminated (**Fig 5G**). Interestingly, the truncated M129 Type-1 still maintained certain stability, suggesting that the A117-A133 interactions were not essential for the stability. In the β-arch, V121-M129 interactions were modest and V121-L125 seemed to take the major role in reinforcing the conformation (**Fig 5H**), while the truncated V129 Type-2 model, in which V122-V129 interactions were the most intense (**Suppl Fig S4B**). The apparent instability of the truncated Type-2 model than the Type-1 counterpart implied significance of A117-A133 interactions for the Type-2 model in the initial phase of conversion. Although we could not find substantial advantages of A133V mutation for structural stability of the completely-refolded Type-2 model [17], V133 might precipitate the β-arch formation in the very initial phase of refolding. This effect may contribute to facilitated transmission of certain scrapie strains by the equivalent polymorphism at codon 136 of ovine PrP [33].

## Discussion

### Relevance of the present/Type-1 model as a local structural model of PrP^Sc^

The present/Type-1 local structural model of PrP^Sc^ provides an example of the structure that is more stable with M129 than V129 or L129, due to its sensitivity to the length of hydrophobic side chain of codon 129. The destabilizing effects of L129 may be associated with the lower susceptibility of elk with L132 polymorphism to CWD [9][10]. Our models may also address the persistency of traits of V129-compatible PrP^Sc^ even after passages to M129-PrP-expressing hosts, showing no abbreviation in incubation on the secondary transmissions and swiftly resuming the original phenotypes with 19-kDa core on back-transmissions to V129-expressing hosts [19][20][21]. If M129-compatible and V129-compatible PrP^Sc^ have β-sheet(120-122) in a relation of opposition in terms of directions of the side chains as in our models, they are not easily interchangeable and may explain the persistence. This can also provide a clue to the mechanism of NAPA [22]. Moreover, we noticed that a final product of coarse-grain MD simulation of PrP-derived peptide encompassing 120-144 had a similar structure as the U-shaped β-arch of our model [34]. Those support the relevance of our models.

### Significance of codon 129 polymorphism on local structures and entire conformations of PrP^Sc^

The Type-1 and Type-2 models/β-arches were rather differently affected by the M/V129 polymorphism and the N- and/or C-terminal truncations. The higher stability of the CN-truncated M129 Type-1 than that of CN-truncated V129 Type-2 β-arch (**Fig 4G vs 5C**) was unexpected. We had anticipated the opposite, because heterozygous CJD cases tend to exhibit more features of V129-homozygous CJD, with 19-kDa core and plaque-type depositions [5][35], suggesting a possibility that V129-PrP predominates the phenotype because it is more amyloid-prone. On the other hand, the higher stability of the M129 β-arch seemed consistent with the higher relative risk of sporadic CJD in M129-homozygous population than the V129-homozygous [36]. The stability of the M129 Type-1 β-arch is presumably attributable to the backbone flexibility due to the absence of Cβ branching and long hydrophobic side chain of M129. It adapted to the C- and/or N-truncations, by modulating the interaction network patterns as if compensating for the lost interactions, interestingly with V121-M129 interactions keeping almost the same moderate proximity scores at ~200 (**Fig 4J**). Possibly, the structural flexibility of M129 Type-1 β-arch that efficiently adapts to the surrounding situations makes it compatible with various conformations of PrP^Sc^ including human-mouse chimeric PrP [8][37], fatal familial insomnia caused by D178N mutation [38], and new-variant CJD [6][7]. The moderate V121-M129 interactions may be important for the stability, because they would not restrain dynamic behaviors of β-sheet(120-122) and β-sheet(128-133). Excessively intense interactions can hamper adaptive changes in their behaviors and futilely preserved the old interaction network, like V121-V129 interactions of the C-truncated V129 Type-1 (**Fig 4E**). The accentuated instability of the C-truncated M129V amyloid suggests that the influences of V129 are greater in the earlier phase before completion of the local conversion (**Fig 4C**; **Fig 6**).

**Fig 6.**
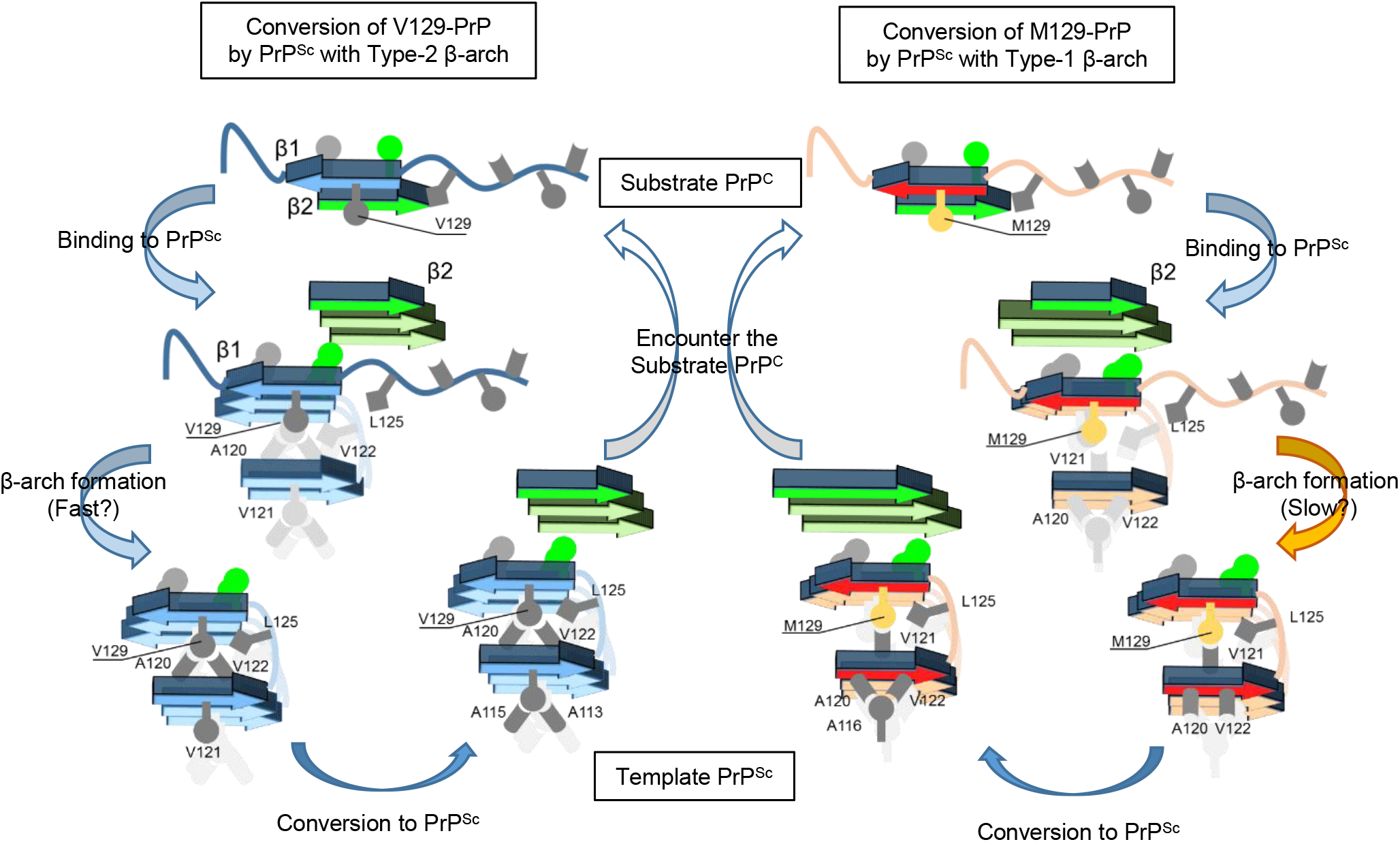
PrP^C^-PrP^Sc^ conversion reactions by two PrP^Sc^ with different types of U-shaped β-arches. A schematic illustration of the PrP^C^-PrP^Sc^ conversion reactions focusing on the U-shaped β-arch region: the conversion of V129-PrP by a PrP^Sc^ with V129 Type-2 β-arch (left half) and the conversion of M129-PrP by a PrP^Sc^ with M129 Type-1 β-arch (right half). The conversion reaction is assumed to begin with PrP^C^-PrP^Sc^ binding, where the template PrP^Sc^ presumably breaks the native anti-parallel β-sheets of PrP^C^, β1 (encompassing 128-131; blue or red arrow) and β2 (encompassing 161-164; green arrow), and incorporates each of them into the corresponding region of the template PrP^Sc^. Subsequently, U-shaped β-arch formation by β1 follows, but this process may be relatively inefficient in the M129 Type-1 β-arch formation due to its back-bone flexibility and the long side chain. Meanwhile, another refolding process may initiate at the β-sheet incorporating the β2 (green stack). Note that β-sheet(120-122) has the 'more-hydrophobic’ side with two hydrophobic side chains of A120 and V122 and the less-hydrophobic side. As the two types of β-arches have the β-sheet(120-122) in a relation of opposition in terms of the direction of the more-hydrophobic side, they would not be easily interchangeable.

Unlike M129 Type-1, the V129 Type-2 β-arch was sensitive to the truncations. The interaction network patterns were characteristically almost the same among full-length and truncated molecules (**Fig 5E**), possibly because the intense bond between β-sheet(120-122) and β-sheet(128-133), i.e., A120-V129 and V122-V129, disabled their adaptation to changes in surroundings. That may eventually destabilize amyloids unless the conformation is preferable for the β-arch, as in the C-truncated V129 Type-1 model (**Fig 4B**). How can a PrP^Sc^ adopt such an intolerant structure? We hypothesize that a refolding pattern initiated at the β-arch can produce PrP^Sc^ with V129 Type-2 β-arch. Hypothetically, the PrP^C^-PrP^Sc^ conversion initiates with incorporation of β1 or β2 (encompassing 128-131 and 161-164, respectively) into the template PrP^Sc^, followed by U-shaped β-arch formation by β-sheet(120-122) and β-sheet(128-130) (**Fig 6**). In this step, the nascent β-sheet(120-122) may preferentially direct hydrophobic side chains of A120 and V122 toward V129, forming the Type-2 β-arch, because it can conceal more hydrophobicity from water and is thermodynamically favorable. Then, the refolding propagates thereof, forming interaction networks in favored ways for the V129 Type-2 β-arch. Thus, as the U-shaped β-arch with V129 is permissive only to preferred local structures, V129-PrP may propagate a limited variety of prion strains. The hypothesis may explain why the Tg mice expressing human-mouse chimeric V129-PrP were resistant even to V129-homozygous CJD [8][37], as discussed in more detail below, and also the resistance of V129-PrP to variant CJD.

Besides structural flexibility, rates of local refolding can affect PrP^Sc^ conformation in theory. Although completely-refolded M129 Type-1 β-arch is stable, the refolding may not be necessarily efficient because of steric effects of the long side chain and the back-bone flexibility of methionine (**Fig 6**), as suggested by relatively slow in vitro aggregation of Y145Stop mutant of Syrian hamster PrP [39] and slower aggregation rates of PrP(120-144) peptide with M129 than that with V129 [40]. If the local refolding around M129 is delayed, refolding processes initiated at another region may propagate meanwhile and eventually predominantly determine the conformation of the entire molecule. In summary, V129 seems to effectively influence conformation determination of PrP^Sc^ by being permissive only to its favored ones, while M129 is apparently subordinate due to its adaptivity and kinetics.

### Other conversion-initiation sites and conformation determination of PrP^Sc^

In the above discussions, we assumed that there are multiple sites where PrP^C^-PrP^Sc^ conversion processes initiate and each prion strain uses its preferred site to propagate refolding in its unique way, resulting in the strain-specific conformation, as we previously hypothesized [29]. We consider the region comprising β1 as one of the sites, and there should be more. For example, P102L mutation of human PrP that causes an inherited prion disease, Gerstmann-Streussler-Sheinker syndrome (GSS) [41][42][43], and I109 polymorphism of bank vole PrP that causes spontaneous transmissible prion disease [44] suggest that the region comprising codon 102 or 109 can autonomously maintain amyloid structures and propagate refolding. To note, the majority of P102L-associated GSS patients have P102L and M129 on the same allele [45][41], whereas A117V-associated GSS has exclusively V129 [42]. When the refolding propagating from P102L arrives at codon 129, M129 may tolerate the imposed conformation by forming an adaptive β-arch, whereas V129 β-arch disfavors it. A117V is interesting because it almost flanks the U-shaped β-arch. It might favor initiation of refolding from the β-arch or closer positioning of the two β-sheets of the β-arch for stability of the amyloid structures.

On the C-terminal side, the region between first and second α-helices (H1~H2) comprising another native β-sheet of PrP^C^, β2, is a site for PrP^C^-PrP^Sc^ binding [46] and might be a site of initiation of refolding. Cooperation with the distal H3 region, possibly by cross-β-spine formation, may further facilitate binding the template PrP^Sc^ and initiate refolding thereof, as suggested by the transmission experiments of CJD to Tg mice expressing human-mouse chimeric PrP that has mouse sequence in the distal H3 [47] and disulfide-crosslink experiments between H1~H2 and distal H3 [48]. As mentioned above, M129 may be compatible with the conformation imposed by the refolding pattern initiated in the C-terminal site, whereas V129 is not [8][37][49]. Their different compatibilities with the pattern are implied by the N-terminally truncated protease-resistant fragments of 12-13 kDa from MM1- and VV2-CJD [50][51]: the former showed 13-kDa fragment truncated at codon 154 that contained β2, while the latter had shorter fragment truncated at 162 that barely contained β2. If the truncated fragments represent half-converted molecules, the refolding process initiated at β2 propagated N-terminally in the PrP^Sc^ of the CJD with M129, i.e., M129 originally fits in the refolding pattern, whereas the pattern of refolding would not occur in the CJD with V129 and is alien to V129.

### Potential application to other in-register parallel β-sheet amyloids

We demonstrated that the short-MD simulations and interaction network analysis could characterize the influences of mutations that are known to affect transmission efficiencies of prions without a supercomputer. That proved usefulness of this method in investigation of amyloids, and further support that PrP^Sc^ is an in-register parallel β-sheet amyloid. Importantly, our models were designed with the main focus on the U-shaped β-arch, and the other regions were arbitrarily designed for the regional stability without extrinsic reinforcement. Therefore, those structures may differ in the full-length PrP^Sc^. Nonetheless, our models have implications about dynamic behaviors of in-register parallel β-sheet amyloids with U-shaped β-arches. Our studies demonstrated that influences of hydrophobic residues on β-arches depend on the conformation of the β-arches and the interactions with other residues in the hydrophobic cores. Moreover, flexibility of each β-arch (not the back bone) and balances among the β-arches are important for stability of an amyloid molecule; excessively intense bonds between a specific pair of β-sheets can be destabilizing. As U-shaped β-arches are essential for efficient cross-β structure formation, the notion from the studies can contribute to investigation of various amyloids and advance our understanding of their prion-like properties and pathogenesis. Targeting U-shaped β-arches for therapies is worth trying: their stable structures are suitable for structure-based drug design.

## Materials and Methods

Detailed protocols of the short MD simulation and various analyses were as described in the preprint on bioRxiv [17]. Here we briefly describe essential points.

### Modeling PrP(107-143) amyloids and MD simulations

The reasons we focused on this region and decided the N- and C-terminal ends and how we designed the amyloid model were previously described elsewhere [17], except that the β-sheet restraints were set for the residues N108-M109, A115-A116, A120-V122, Y128-L130, P137-F141.

### MD simulations

The protocol for the 10-ns MD simulation was the same as reported elsewhere [17], except that the restraints were slowly removed first in the directions along the X-Y plane, and then in the Z direction (stacking direction).

### Analyses

The protocols for determination of the secondary structure contents, evaluation of hydrophobic contacts, definition of proximity scores, and the method for visualization of the results are the same as previously reported [17]. For evaluation of hydrophobic interactions, we used “proximity score” that indicates persistence of hydrophobic contact, which is defined by distance between centers of mass of sidechains of two given residues less than 5Å [17][32], because it conveniently enables evaluation of intra- and inter-layer interactions at the same time.

### Statistical analyses

Two-tailed Welch two sample t-test was used for analysis of statistical differences between two groups. A p-value less than 0.05 was considered statistically significant.

## Acknowledgement

The numerical calculations were carried out on the TSUBAME2.5/3.0 supercomputer at the Tokyo Institute of Technology and the Reedbush-U supercomputer at the Information Technology Center, the University of Tokyo. This work was supported by the "TSUBAME Encouragement Program for Young/Female Users" of Global Scientific Information and Computing Center at the Tokyo Institute of Technology, the “Initiative on Promotion of Supercomputing for Young or Women Researchers” from the Information Technology Center, the University of Tokyo, Takeda Science Foundation (www.takeda-sci.or.jp/), and JSPS KAKENHI Grant Numbers 19K16058 (to HO) and 19K22539 (to YT).

