## Supplemental Fig S1-4 for "Molecular dynamics simulation of local structural models of PrP^Sc^ reveals how codon 129 polymorphism affects propagation of PrP^Sc^"

### Supplementary Figure S1

Final shots of the ten independent runs of 10-ns MD simulations of wild-type or mutants of the M129 Type-1 model are presented.

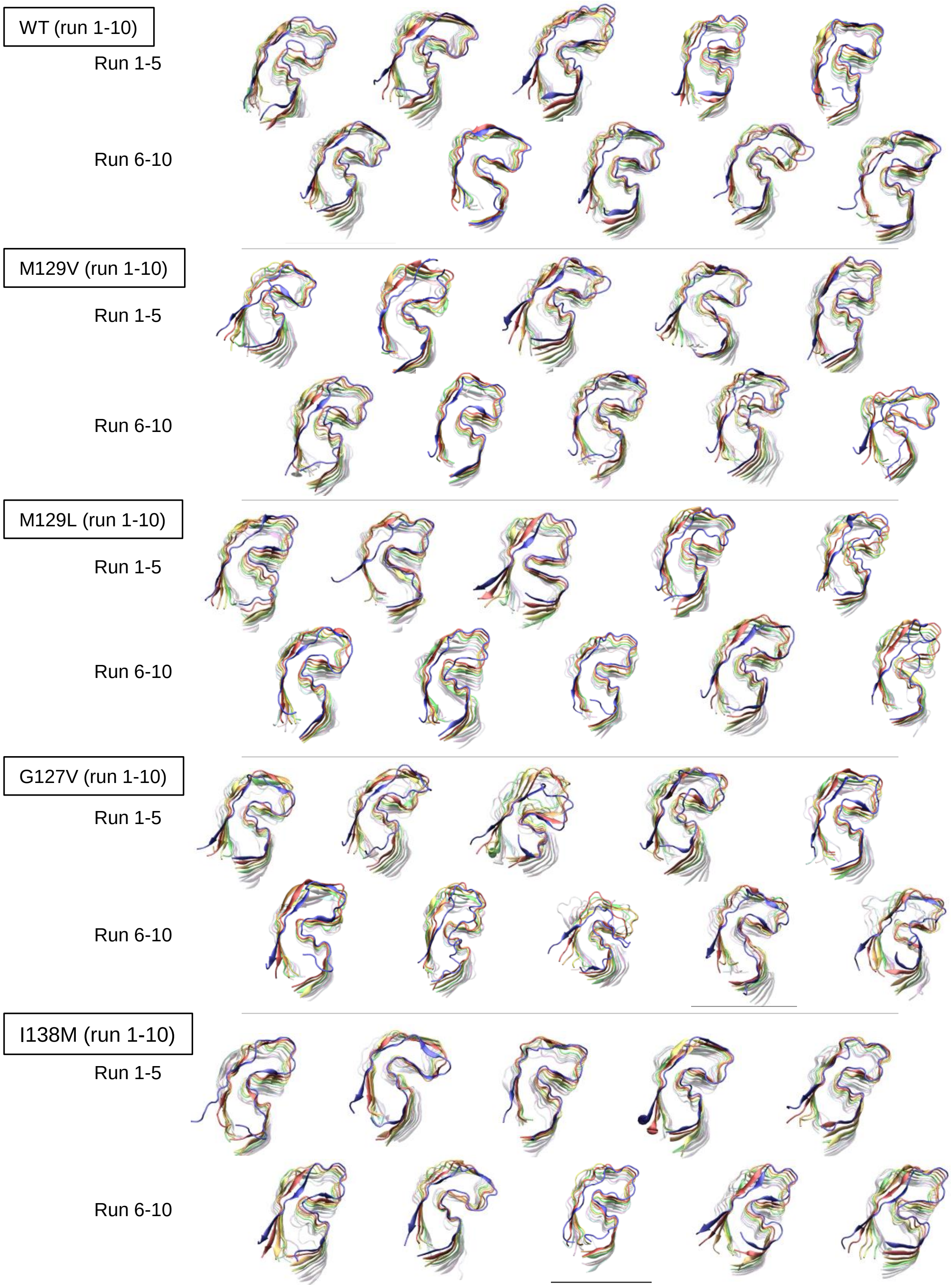

### Supplementary Figure S2

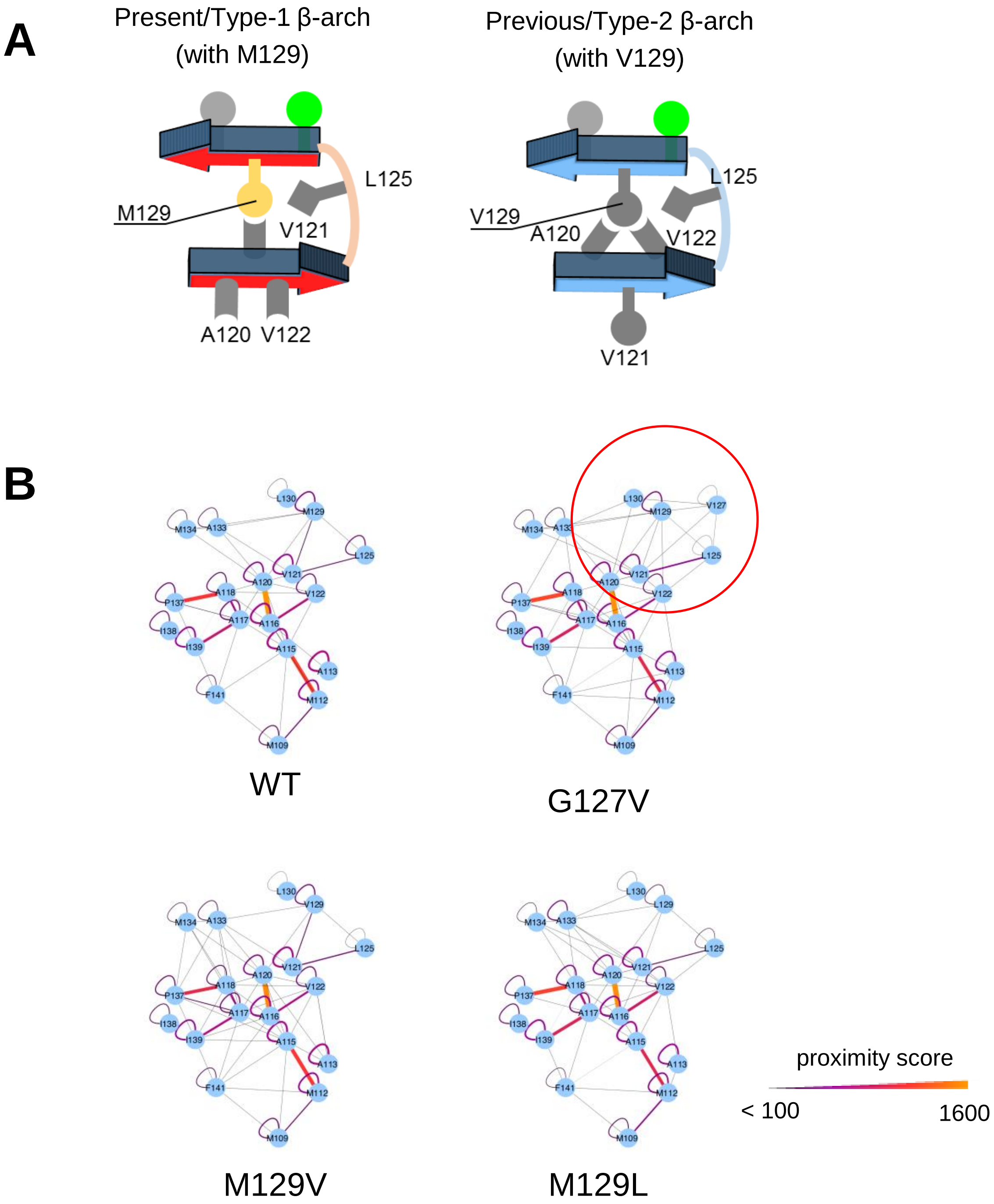


**A.** Schematic illustration of the ‘Type-1’ and ‘Type-2’ β-arches/models (left and right, respectively). The types are defined based on the number of residues that mainly interact with the residue 129. Note that only V121 confronts the residue 129 in Type-1 β-arch, whereas A120 and V122 confront it in Type-2 β-arch.

**B.** Diagrams of hydrophobic interaction networks of the wild-type and mutants of the present models (M129 Type-1 model). Threshold for hydrophobic contact: 5.0 Å.

The colors and thickness of the lines represent the proximity scores for the interactions. The red circle indicates the residues of the U-shaped β-arch.

### Supplementary Figure S3

Final shots of the five independent runs of 10-ns MD simulations of wild-type or mutants of M129 Type-1 and V129 Type-2 models are presented.

C-truncated WT of the present model (M129 Type-1) (run 1-5)

Run 1-5

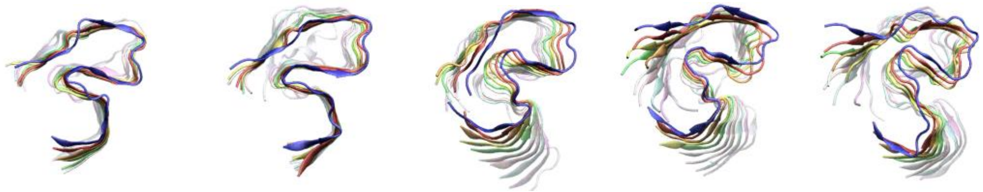

C-truncated M129V of the present model (V129 Type-1) (run 1-5)

Run 1-5

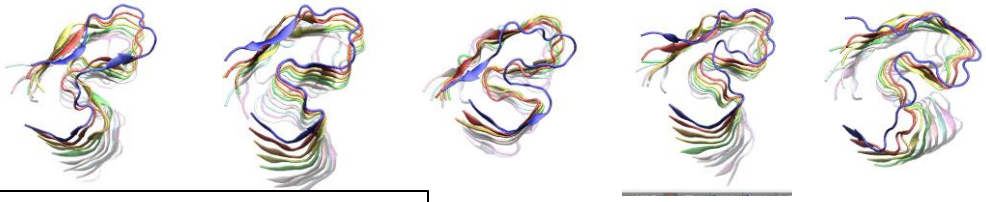

N-truncated WT of the present model (M129 Type-1) (run 1-5)

Run 1-5

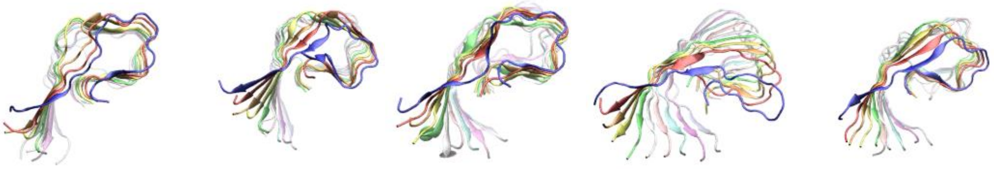

C, N-truncated WT of the present model (M129 Type-1) (run 1-5)

Run 1-5

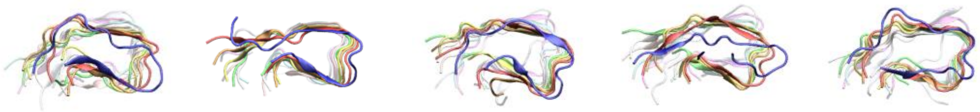

C-truncated M129V of the previous model (V129 Type-2) (run 1-5)

Run 1-5

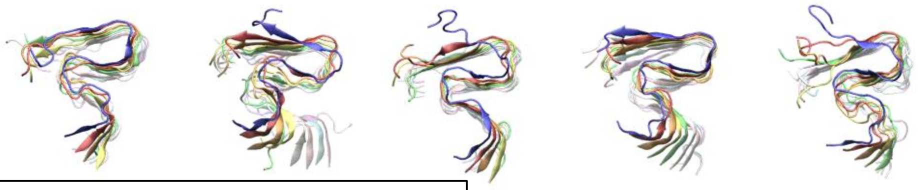

N-truncated M129V of the previous model (V129 Type-2) (run 1-5)

Run 1-5

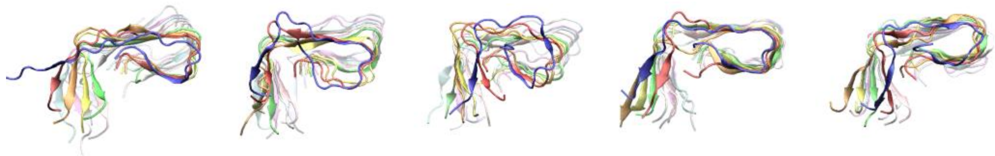

C, N-truncated M129V of the previous model (V129 Type-2) (run 1-5)

Run 1-5

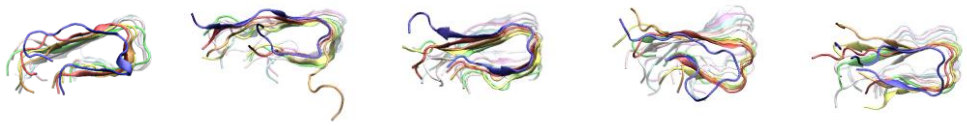

### Supplementary Figure S4

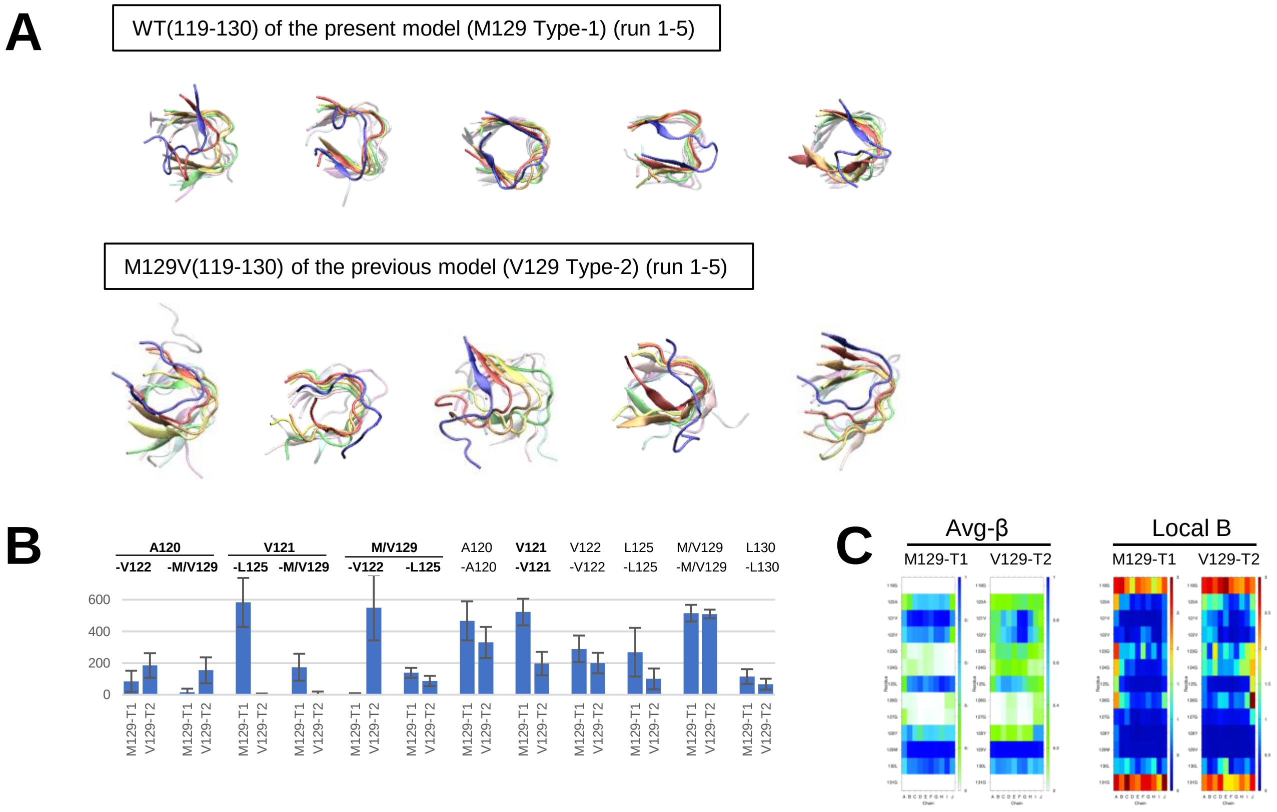


**MD simulations of the peptides corresponding to the residues 119-130 of M129 Type-1 and V129 Type-2 models.**

**A.** Final shots of the five independent runs of 10-ns MD simulations of the peptides.

**B.** A graph of proximity scores for representative hydrophobic interactions. The interactions in bold letters indicate those in the hydrophobic core of the U-shaped β-arch. The bars and error bars represent mean ± SD of scores from five independent runs of 10-ns simulation. M129-T1 and V129-T2, peptides 119-130 of the M129 Type-1 and V129 Type-2 models, respectively.

**C.** Heat maps of Avg-β values and local B-factor values of the peptides based on the five independent runs.
